# p53-Bad* in a Hepatocellular Carcinoma Mouse Model

**DOI:** 10.1101/2023.06.29.547129

**Authors:** Katherine Redd Bowman, Phong Lu, Carol Lim

## Abstract

Recent advances in liver cancer treatments have not changed the fact that the majority of patients will not survive the disease. In order to advance future liver cancer treatments, this work presents an exploration of various iterations of the liver cancer specific AFP promoter as well as the gene construct p53-Bad*. p53-Bad* is a mitochondrially targeted re-engineered p53 therapy that has shown previous success in a zebrafish HCC model. Both the most promising AFP promoter and p53-Bad* were packaged in an adenoviral delivery system and tested in vitro in liver cancer cell lines. Finally, mixed results for adenoviral p53-Bad* in vivo are presented, and this work suggests future modifications to study parameters in order to further explore the potential of p53-Bad* as a potential liver cancer therapeutic.

## Introduction

Hepatocellular carcinoma, or HCC, is the most common type of liver cancer and the third leading cause of cancer death globally^1^. Though survival rates have improved slightly in the past several decades, they remain low—as little as 2.5% in the United States^2^—for advanced HCC. Despite approval of several new systemic treatment regimens in the last few years (e.g., atezolizumab plus bevacizumab^1^ and tremelimumab plus durvalumab^3^) marginally improving the overall survival of patients, the majority of patients with advanced HCC will still succumb to their disease^1^. Alternative therapies are still desperately needed, particularly for patients with advanced HCC.

We have previously reported that p53-Bad*is a re-engineered construct that combines the tumor suppressor p53 with the anti-apoptotic BH3 protein Bad^4-5^. The star (*) indicates two serine to alanine mutations (S112A and S136A) that increase the mitochondrial localization of Bad. In fact, these mutations are enough to overcome the nuclear localization signal of p53, bringing the p53-Bad*construct to the mitochondria, where p53 can induce rapid apoptosis^4-5^. p53-Bad * has shown high efficacy in inducing apoptosis in vitro in four different liver cancer cell lines with varying p53 mutation statuses^4^. Strikingly, p53-Bad*also reduced tumor burden and induced apoptosis in a zebrafish model^4^.

Typically only expressed in the fetal liver and placenta^6^, alpha fetoprotein (AFP) levels are abnormally high in HCC—the protein is expressed in up to 80% of HCC cases^7^. High AFP levels in HCC also correlate with larger tumors, more advanced staging, and decreased survival^8^. AFP promoters have been used in a variety of HCC-specific functions, including AFP-targeted imaging^9^, RNA interference for HCC^10^, and HCC-specific oncolytic adenovirus^11^. Thus, expressing p53-Bad* gene therapy under control of the AFP promoter could not only be efficacious, but also apply to those patients in most need of new therapies.

This work sought to explore the use of p53-Bad* in a mouse model of HCC, as well as the efficacy of different iterations of the AFP promoter. These include the proximal AFP promoter as well as two iterations of the promoter, AFP-E^12^ and AAB-AFP^11^—which include the proximal promoter and variations of the enhancer region. These promoters were tested in liver cancer cell lines first. We then combined the most promising promoter with p53-Bad* in an adenoviral delivery system and tested it in vitro before exploring the efficacy of adenoviral p53-Bad* in vivo in a mammalian mouse model.

## Materials and Methods

### Promoter Cloning

Restriction enzyme cloning was used to create the various alpha-fetoprotein (AFP) promoters discussed above, with PCR run on genomic DNA previously extracted from normal fibroblasts as before^13^ with primer sets to create inserts as follows. pAFP-EGFP: F:5’-GCAGCTGCTAGCCGAATCCGATCGAATATTCTGTAGTTTGAGGAG-3’ and R:5’-GCAGCTACCGGTCGAATCACTAGTTGTTATTGGCAGTGGTGG-3’ pAFP-EGFP-p53-Bad*: F:5’-GTATATGCTAGCATAATCCTCGAGAATATTCTGTAGTTTGAGGAG-3’ and R:5’-GTATATACCGGTATAATTGAATTCTGTTATTGGCAGTGGTGG-3’ pAFPE-EGFP and pAFPE-p53-Bad*: GFP F:5’-GCACGGACCGGTTTATATATCAAAATAAACTTGAG-3’ p53-Bad* F:5’-GCCGCGGAATTCTTATATATCAAAATAAACTTGAG-3’ R:5’-CTATTAACCGGTTAGGAAGTTTTCGCAATAATAC-3’ pAAB-AFP-EGFP and pAAB-AFP-p53-Bad*(3 inserts): (1) F:5’-GATACTGCTAGCCAGATTGAATTATTTGCCTG-3’ R:5’-GATACTCAATTGTTGTAGCCATGTGAGTGG-3’ (2) F:5’-GATACGCAATTGCAGATTGAATTATTTGCCTG-3’ R:5’-GATACTTGTACATTGTAGCCATGTGAGTGG-3’ (3) F:5’-GATACTTGTACACCTGATTAATAATTACACTAAG-3’ GFP R:5’-CTTATACGATCGTAGGAAGTTTTCGCAATAATAC-3’ p53-Bad* R:5’-CTTATACTCGAGTAGGAAGTTTTCGCAATAATAC-3’.

### Cell Lines and Maintenance

All cell lines were purchased from ATCC (Manassas, VA) and grown in flasks in monolayers at 37°C with 5% CO_2_. Four liver cancer cell lines were used, including HepG2/C3A (C3A) (derivative of HepG2, p53 WT), Hep3B2.1-7 (Hep3B, p53 null), PLC/PRF/5 (p53 dominant negative R249S mutation), and Snu398 (p53 null), as well as four normal cell lines, THLE-2 (immortalized normal liver cells), hFOB1.19 (immortalized normal bone cells), IMR-90 (normal fetal lung cells), and BJ (normal foreskin fibroblasts). Cells were maintained in DMEM (C3A, Hep3B2.1-7, Snu398, PLC/PRF/5, and IMR-90), RPMI 1640 (Snu398), DMEM/F12 (hFOB1.19), or BEGM (THLE-2). Media was supplemented with 1% L-glutamine (Corning), 1% penicillin-streptomycin (Gibco), and 10% FBS (Atlanta Biologicals). THLE-2 cell media was also supplemented with 5 ng/mL epidermal growth factor (Corning) and 70 ng/mL phosphoethanolamine (Sigma-Aldrich), and the gentamycin/amphotericin and epinephrine provided with the BEGM media kit (Lonza, Walkersville, MD) were discarded as per ATCC instructions. THLE-2 cells were grown in flasks coated in fibronectin (Sigma-Aldrich), bovine serum albumin (Sigma-Aldrich), and bovine collagen (Advanced Biomatrix) as per ATCC instructions. All experiments were conducted between cell passages 5 and 20.

### GFP Assay

Cells were seeded at 500,000/well for THLE-2, IMR-90, hFOB1.19, and BJ and 400,000/well for C3A, PLC/PRF/5, Snu398, and Hep3B2.1-7 in 6-well CellBIND plates (Sigma Aldrich) for GFP assays. After 24 hours, each well was transfected with 1 pmol of DNA using JetPrime transfection reagent (PolyPlus Transfection) according to the manufacturer’s instructions. 48 hours post-transfection, cells were collected and treated as before^13^. In brief, cells were pelleted and resuspended in 300 μL PBS, then analyzed for GFP via flow cytometry (FACS-Canto-II with FACS Diva software, Core Facility, University of Utah)^13^. Experiments were repeated in triplicate for each gene construct 3 times, with a representative experiment for each cell line shown without normalization to display raw transfection numbers, and normalization to UT (set as 0%) and CMV-GFP (set as 100%) performed to combine data for comparison between cell lines. 2-way ANOVA with Tukey’s post-test was run in GraphPad Prism 9.4.1 (multiple comparisons not shown in figure).

### Adenoviral Vectors

Empty adenovirus (Ad), adenoviral WT p53 (Ad-p53-wt), adenoviral p53-Bad* (Ad-p53-Bad*), adenoviral AAB-AFP-p53-Bad* (Ad-AAB-AFP-p53-Bad*), and adenoviral Bad* (Ad-Bad*) were produced by SignaGen Laboratories (Maryland, USA). All adenoviruses are based on recombinant serotype 5 (Cat# SL100702, Ad.MAX System with E1/E3 deletion). The final purified virus titers were determined to be 1.57×10^11^ pfu/mL, 5.68×10^10^ pfu/mL, 1.04×10^11^ pfu/mL, 1.26×10^11^ pfu/mL, and 5.05×10^10^ pfu/mL respectively. The adenoviruses were used in all MTS assay and mouse study figures. They will be referred to in the text and figure legends simply as empty (or null), WT p53, p53-Bad*, AAB-AFP-p53-Bad*, and Bad*, respectively.

### MTS Assays

Cells were seeded 10,000 per well into a transparent 96 well plate, then allowed to incubate for 24 hours before treatment with viral constructs diluted to the appropriate MOI in cell culture medium. After incubation for 48 or 72 hours, cells were treated with CellTiter 96® AQueous One Solution Cell Proliferation Assay reagent (Promega) according to the manufacturer’s instructions. Aer incubation for 1-4 hours, cell plates were analyzed for absorbance at 490 nm using a Cytation 3 imaging reader (BioTek Instruments). Data was plotted and analyzed by one-way ANOVA with Tukey’s post-test in GraphPad Prism 9.4.1.

### Mouse Studies

6-7 week old female PRR NRG mice were subcutaneously injected with 1×10^6^ Hep3B cells, after which tumors were allowed to grow. Tumors were measured and volumes calculated, and 5 mice per group were enrolled in two waves when tumors were 50-250 mm^3^ in volume. Virus was injected intratumorally in doses of 1×10^9^ PFU in 40 uL PBS 3x/week with a total of 8 doses, and mice were euthanized and tumors collected on day 18 (1 day after final dose). All mouse studies were performed at the Preclinical Research Resource at the Huntsman Cancer Institute, study ID: 221101CLim_Hep3B, IACUC: 21-1000. 2-way ANOVA with Tukey’s post test was used to analyze data in GraphPad Prism 9.4.1.

### qRT-PCR

Aer subject termination at the end of the study, mouse liver tumor samples were collected and frozen at -80 °C. These samples were then ground in TriReagent (Zymo Research) and vortexed before RNA extraction using Direct-zol RNA miniprep kits (Zymo Research). All RNA samples were diluted to the same concentration before using the Superscript III First-Strand Synthesis System (Thermo Fisher) to synthesize cDNA according to the manufacturer’s instructions using half oligodT, half random hexamer primers. qRT-PCR was run on a StepOne Plus Real-Time PCR machine (Applied Biosystems) with Taqman Universal PCR Master Mix (Thermo Fisher) and pre-designed Taqman probes targeting 18S (Hs99999901_s1), Bad (Hs00997773_m1), and p53 (Hs01034254_g1TP53). qRT-PCR was run in duplicate for each tumor sample, then normalized with a standard curve and expression of 18S in Microsoft Excel before analysis in GraphPad Prism 9.4.1. One data point in tumor B.L1 (p53-Bad*) was run in duplicate but only one 18S datapoint was used to normalize due to error in the second standardization datapoint. Two data points were removed, one each from the Bad* and WT p53 data sets, after Grubb’s test cleaning (see Figure S2). Both of these data points also had lower than expected 18S (housekeeping gene) expression levels, indicating lower concentration than expected, which can propagate measurement error.

## Results

Several iterations of the AFP promoter—AFP, AFP-E, and AAB-AFP—were tested in four HCC cell lines and four normal non-HCC cell lines (Figure 1). HCC cell lines varied in AFP levels, and included C3A (AFP +++), Hep3B2.1-7 (AFP ++), PLC/PRF/5 (AFP +), and Snu-398 (AFP -). Non-HCC cell lines included THLE-2 (normal liver), IMR-90 (normal lung), hFOB1.19 (normal bone) and BJ (normal foreskin). C3A cells showed fairly high transfection rates with all AFP promoters, with AAB-AFP showing the highest transfection rate of around 40%, around two-thirds the transfection level of the CMV control promoter. Hep3B2.1-7 cells showed some efficacy for the AFP-E and AAB-AFP promoters, though very little for the AFP promoter. AAB-AFP also showed some activity in PLC/PRF/5 and Snu-398 cells, demonstrating around 30 and 25 transfection percent compared to the CMV control promoter. All non-HCC cells showed very low activity for all AFP promoter iterations. Overall, when transfection rates are normalized to CMV as 100%, the AFP promoter displays specificity for C3A cells, AFP-E for C3A and Hep3B2.1-7 cells, and AAB-AFP for all HCC cells, with AAB-AFP showing the highest transfection efficacy.

**Figure 1.**
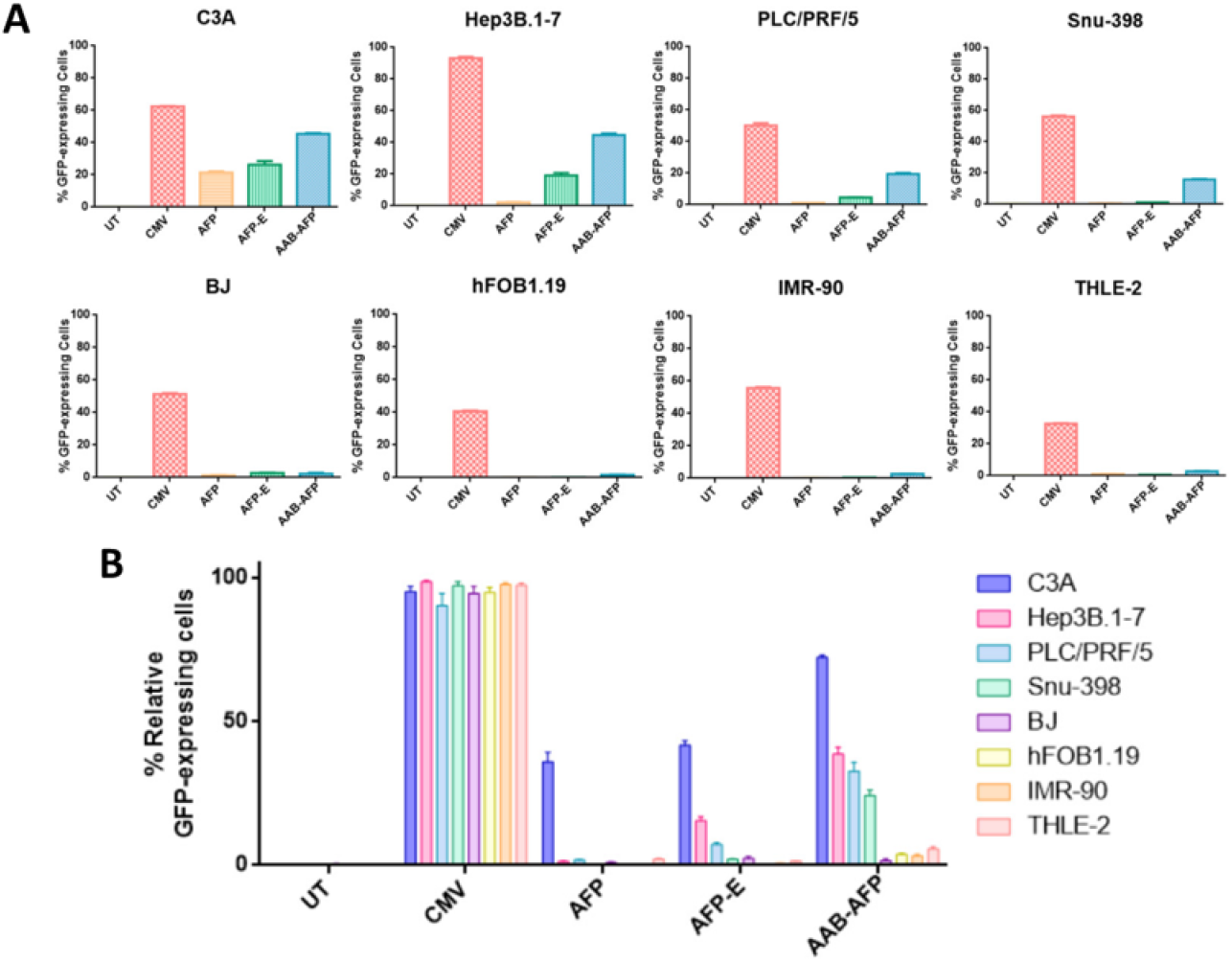
AFP promoters in liver cancer and normal cell lines. Liver cancer (A, top) and normal cell lines (A, bottom) were treated with plasmids encoding EGFP with four different promoters: CMV (constitutive control), AFP, AFP-E, and AAB-AFP. After normalizing to CMV as 100%, data from each cell line was compared (B). 2-way ANOVA with Tukey’s post-test was run for (B) (multiple comparisons not shown on graph).

Aer seeing successful apoptotic activity in HCC cell lines and tumor reduction/apoptosis in a zebrafish liver cancer model^4^, p53-Bad* was packaged in an adenoviral vector for testing in a subcutaneous xenograft mouse model. Before testing the virus in mice, adenoviral vectors null (empty vector), WT p53, p53-Bad*, Bad*, and AAB-AFP-p53-Bad* (most effective tested HCC-specific promoter with p53-Bad*), were used to treat Hep3B cells (p53 null). MTS assays at 48 and 72 hours showed that, contrary to our previous non-viral apoptotic in vitro experiments, WT p53 was the most effective construct at inducing apoptosis at 48 hours post-treatment (Figure 2, top). This effect was no longer significant at 72 hours post-treatment, at which point p53-Bad* and Bad* had no significant difference in efficacy at inducing loss of cell viability in Hep3B cells (Figure 2, bottom). MTS assays were also performed in PLC/PRF/5 liver cancer cells (p53 dominant negative mutation, R249S^4^). Adenoviral p53-Bad* was less effective in PLC/PRF/5 cells (Figure 3), which was unexpected based on our previous apoptotic data in the cell line^4^. One possible cause of adenoviral p53-Bad* inducing less cell death than adenoviral WT p53 may be the fact that p53-Bad* had distinctly lower expression levels compared with WT-p53 (and Bad*, see figure S1) in the adenoviral system. Due to the success of adenoviral p53-Bad* at causing extensive cell death at 72 hours in Hep3B cells (the planned cell line for the mouse model), we proceeded with mouse studies. Adenoviral AAB-AFP-p53-Bad* was excluded from the mouse study because the expression levels of construct were simply not high enough with the viral delivery system to induce cell death.

**Figure 2.**
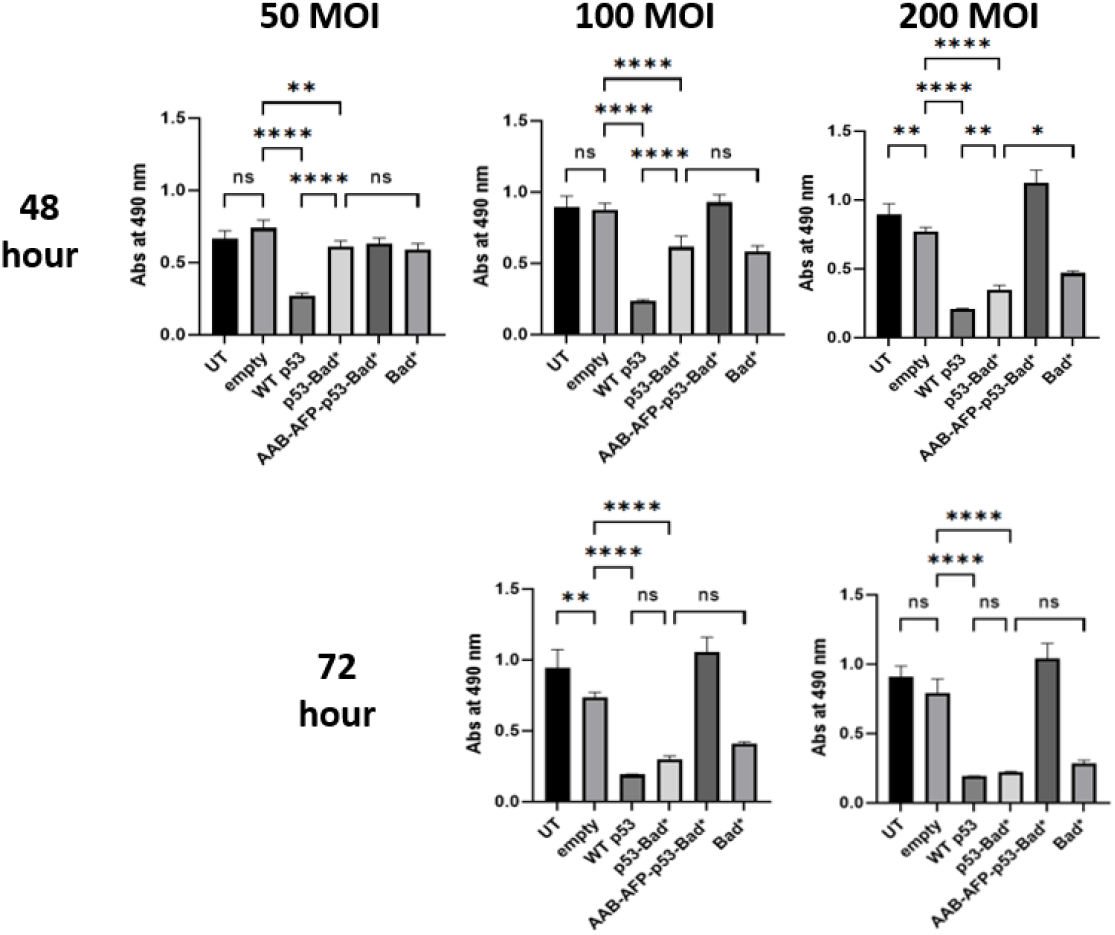
MTS assay for Hep3B cells 48 and 72 hours post viral treatment. Increasing the multiplicity of infection (MOI) improved the efficacy of Bad* and p53Bad* at reducing cell viability at a 48 hour timepoint but had little effect on the efficacy of WT p53. At 72 hours post treatment, there was no significant difference in efficacy between adenoviral WT p53, p53-Bad*, or Bad* at 100 MOI or 200 MOI. Stats: One-way ANOVA with Tukey’s post-test. Ns = not significant, *p<0.05, ^**^p<0.01, ^****^ p<0.001.

**Figure 3.**
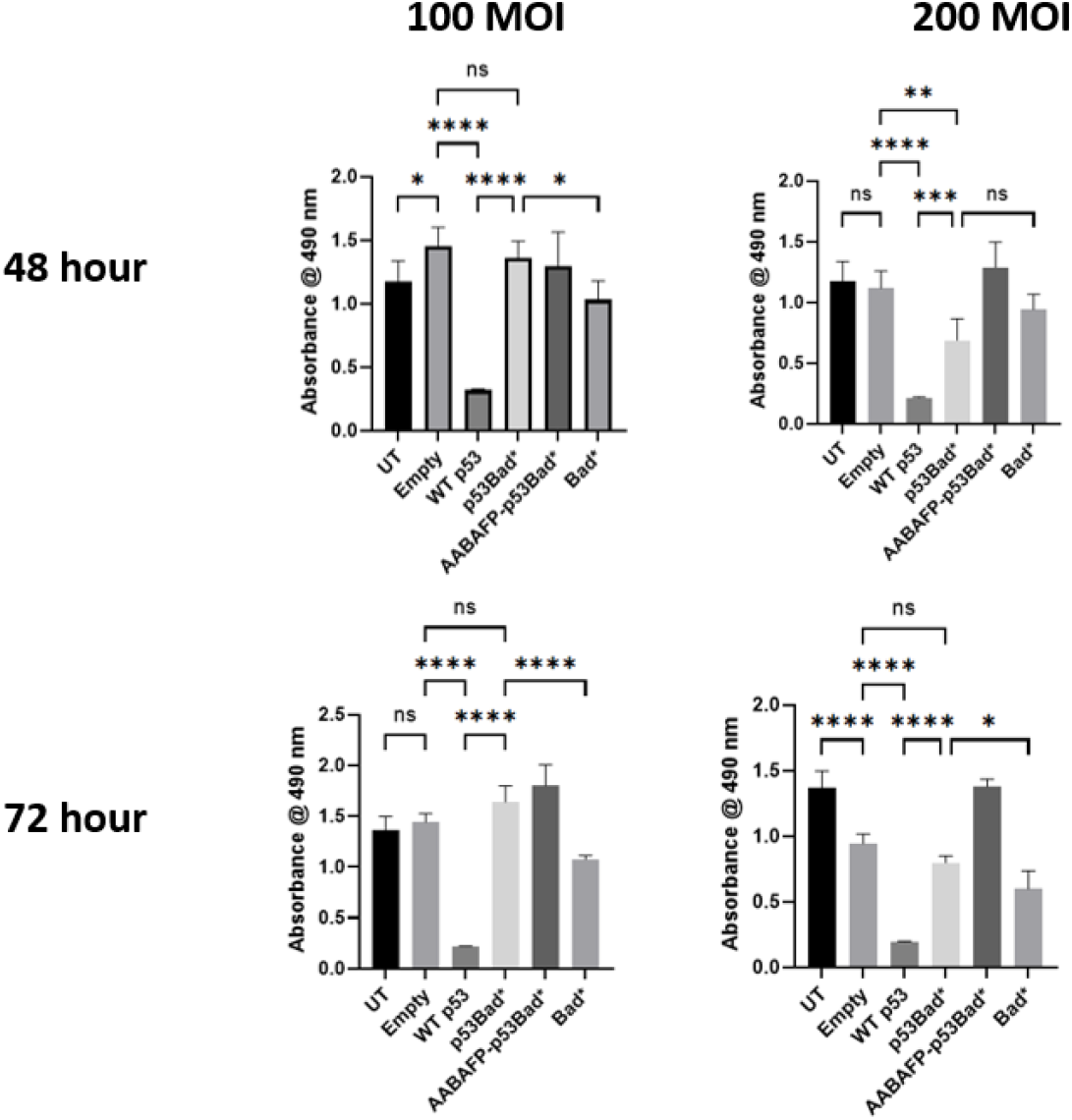
MTS data in PLC/PRF/5 cell line. Ad-WT p53 significantly more effective at reducing cell viability compared to all other constructs at all doses and timepoints. This is consistent with results from apoptosis assays in our previous publication, but the decreased efficacy of Ad-p53-Bad* is not. Stats: one-way ANOVA with Tukey’s post-test, ns=not significant, *p<0.05, ^**^p<0.01, ^***^p<0.001, ^****^p<0.0001.

Mice were treated with PBS (vehicle), null (empty Ad vector), and vectors with adenoviral WT p53, p53-Bad*, or Bad*. Animal weights, when normalized to starting weight, had no significant difference throughout the study (Figure 4, left). Tumor volumes, when normalized to starting volume, had no significant difference until day 14, at which point adenoviral WT p53 and Bad* had significantly smaller volumes compared with PBS-treated tumors (Figure 4, right). By day 18, all adenoviral constructs had significantly smaller volumes compared to PBS-treated tumors, but only WT p53 was also significantly smaller than null (empty adenovirus) treated tumors.

**Figure 4.**
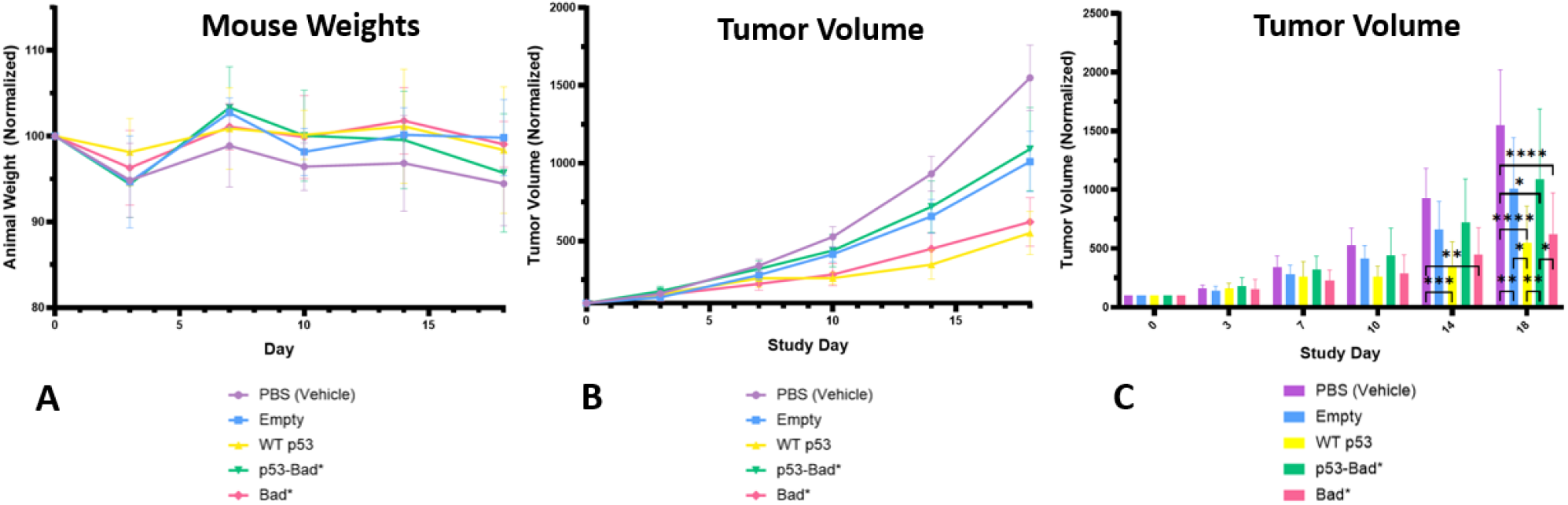
Mouse weight and tumor data. Each group consisted of 5 female PRR NRG mice, with viral doses given 3x per week for a total of 8 doses (final dose on day 17, final measurement on day 18). When normalized to their starting weights, the average weights of animals in each group had no significant difference at any timepoint in the study (A). When normalized to their starting values at enrollment, tumor volumes were not significantly different for any group on days 0, 3, 7, and 10 of the study (B, C). On day 14 of the study, ad-WT p53 treated tumors were significantly smaller than PBS (vehicle) treated tumors. On day 18 of the study, all cither groups were significantly smaller than PBS (vehicle) treated tumors, ad-WT p53 and ad-Bad* treated tumors were significantly smaller than ad-p53-Bad* treated tumors, and ad-WT p53 treated tumors were significantly smaller than empty adenovirus treated tumors. All comparisons not shown were not significant (ns). 2-way ANOVA with Tukey’s post-test was used to analyze both data sets, with ns = not significant, *p<0.05, ^**^p<0.01, ^***^p<0.001, and ^****^p<0.0001.

Because the results of this in vivo study differed from our non-viral data both in vitro and in vivo in zebrafish^4^, we decided to explore potential causes for these results. One major variable in the mouse study was the starting size of the Hep3B tumors, which varied from ∼50-250 mm^3^.

Though tumors that started smaller displayed a wide range of final sizes, tumors that started in the larger range (>110 mm^3^) all finished in the upper 50% range of tumors, with the exception of one WT p53 treated tumor (Figure 5). This suggests that a narrower range of starting tumor size, and smaller starting tumor size, should theoretically improve the quality of the data, and, were this study to be run again, the authors would recommend a tumor starting size range of 50-100 mm^3^.

**Figure 5.**
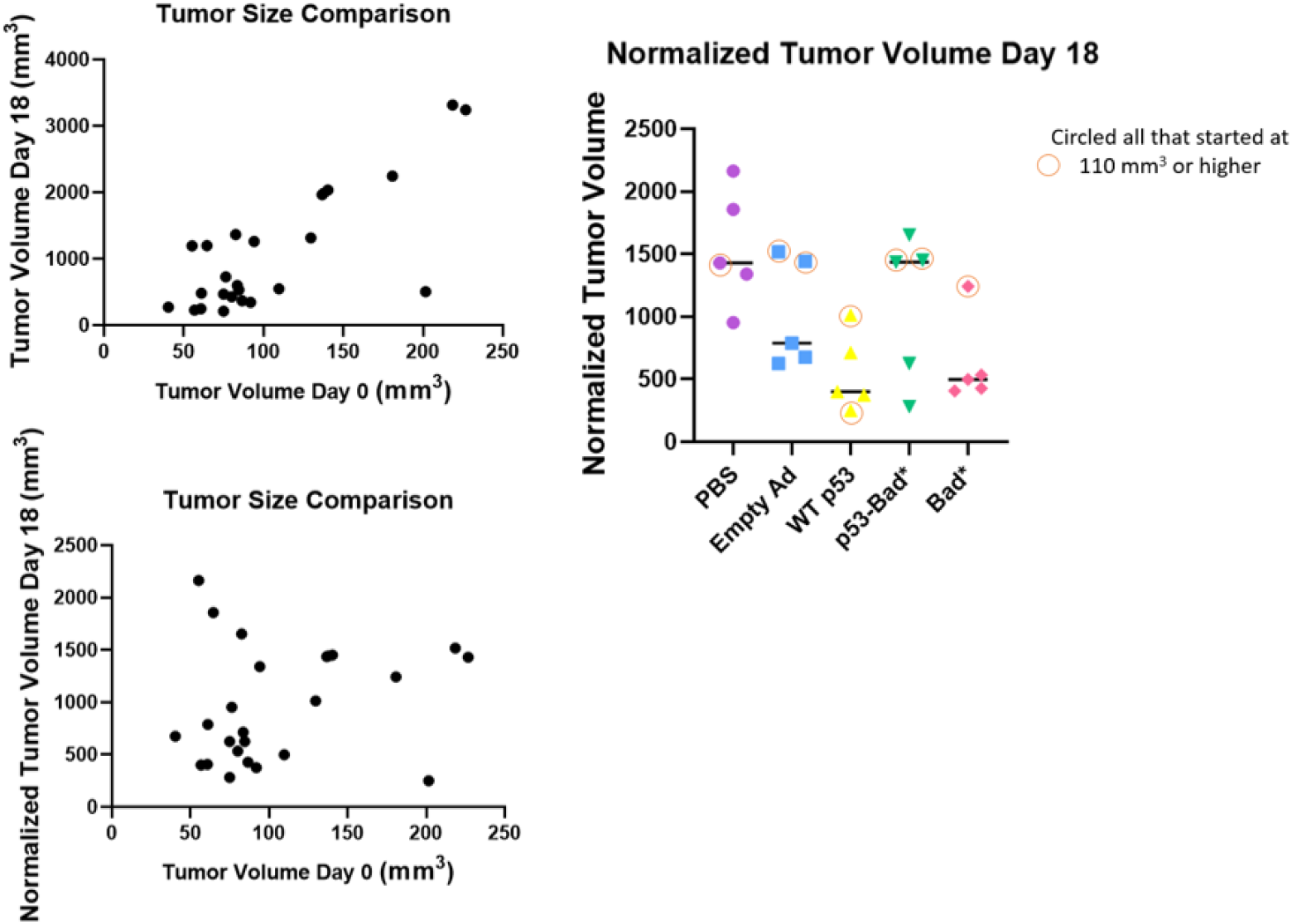
Tumor Size Comparison. Overall tumor growth appears to be affected by the starting size of the Hep3B tumors. With the exception of one ad-WT p53 treated tumor, all tumors that began the study in the upper range of size (> 110 mm3) demonstrated growth in the top 50% of tumors.

Next, qRT-PCR was used to measure the RNA expression levels of human p53 and human Bad present in the adenoviral treated tumors. Adenoviral WT p53 and p53-Bad* would both be expected to express human p53 at similar levels, and Bad*/p53-Bad* would both be expected to express human Bad at similar levels. Interestingly, we found that WT p53 treated tumors expressed significantly more human p53 than p53-Bad* treated tumors, and Bad* treated tumors expressed significantly more human Bad than p53-Bad* treated tumors (Figure 6D).

**Figure 6.**
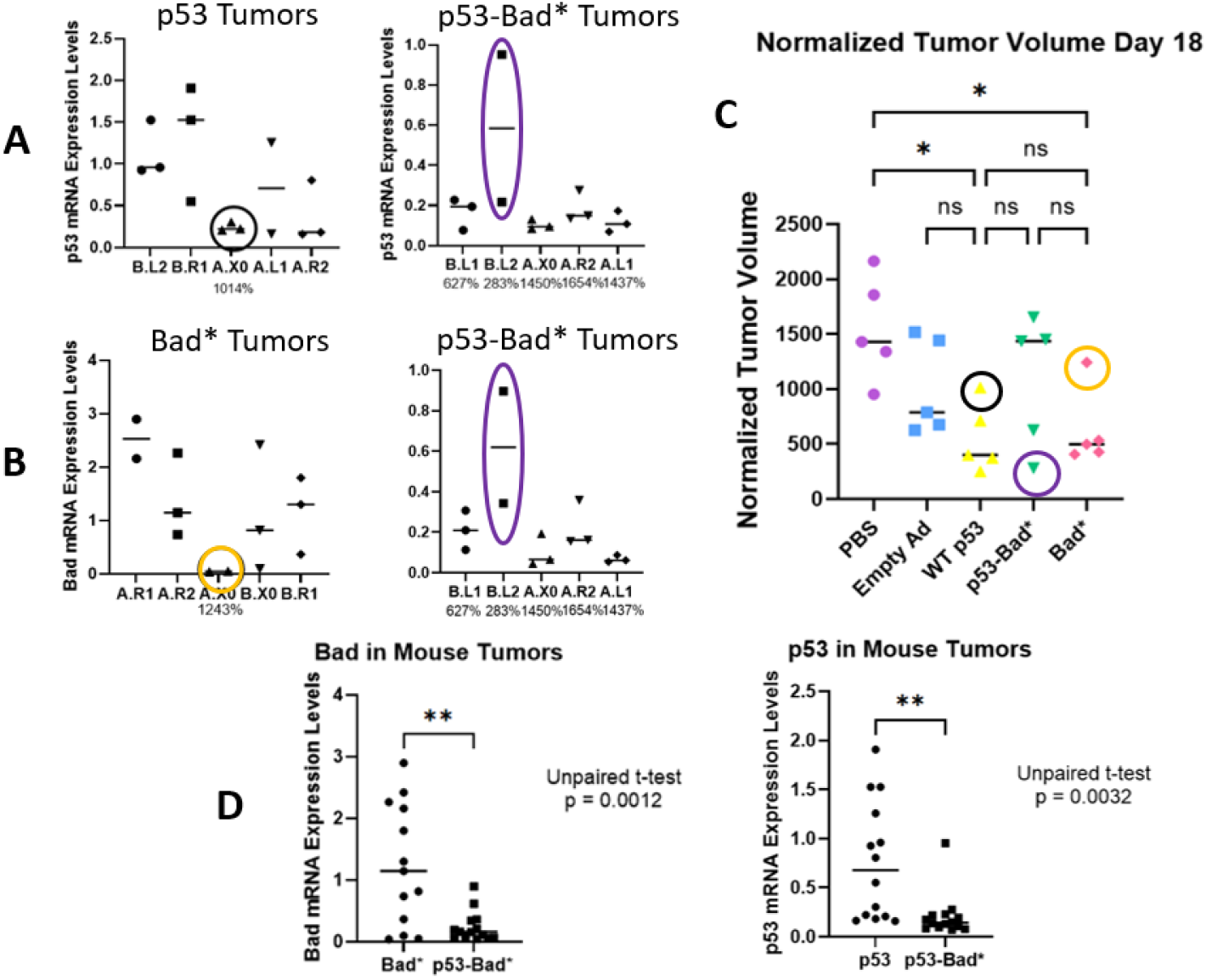
p53 and Bad RNA expression levels in mouse tumor samples. Samples from each treated tumor were analyzed for human p53 RNA (p53 and p53-Bad* tumors, A in the figure) or human Bad RNA (Bad* and p53-Bad* tumors, B in figure) When compared with the normalized tumor volume data from day 18 of the study (C), the ad-p53 treated tumor with particularly low human p53 expression was the largest tumor in its group, the ad-Bad* treated tumor with particularly low human Bad expression was the largest tumor in its group, and the ad-p53-Bad* treated tumor with higher overall expression levels of human p53 and Bad was the smallest tumor in its group. Overall, the ad-Bad* treated tumors had significantly higher human Bad expression than the ad-p53-Bad* treated tumors, and the ad-WT p53 treated tumors had significantly higher human p53 expression than the ad-p53-Bad* treated tumors (D). One outlier each v/as removed from the p53 and Bad* tumor data after running Grubb’s test on the data. (C) was analyzed with One-way ANOVA and Tukey’s post test (ns= not significant, *p<0.05). (D) were analyzed with unpaired t-tests (p values labeled on graphs).

Within groups, we saw a correlation between larger tumors in the WT p53/Bad* categories and low p53/Bad expression (Figure 6A-C), as well as correlation between smaller tumors and higher p53/Bad expression in p53-Bad* treated tumors (Figure 6A-C).

## Discussion

This work tested the proximal AFP promoter as well as two iterations of the promoter, AFP-E ^12^ and AAB-AFP ^11^, which include the proximal promoter and variations of the enhancer region. All three promoters showed efficacy in highly AFP-positive C3A cells, with decreasing transfection levels that correlated with decreasing AFP levels in Hep3B2.1-7, PLC/PRF/5, and Snu398 cells.

Interestingly, the promoter with the highest expression across all HCC cell lines, AAB-AFP, still showed some expression in AFP-negative Snu398 HCC cells (∼25% of the transfection rates of CMV control promoter levels). When tested in normal cells, however, AAB-AFP transfection levels fell drastically (∼1-8% of the CMV control promoter levels, depending on the cell line). This indicates that, even in non-AFP expressing HCC tumors, the AAB-AFP promoter could still be efficacious enough to treat tumors while sparing most normal cells. Normal cell lines tested included normal liver cells (THLE-2), fibroblasts (BJ), lung cells (IMR-90), and bone cells (hFOB1.19). As two of the most common sites of HCC metastases ^14^, lung and bone cells were included to show lack of expression in these cells as an early step in the potential development of p53-Bad* as a systemic HCC treatment. Unfortunately, when used to express p53-Bad* in an adenoviral vector rather than GFP in a plasmid, the efficacy of AAB-AFP dropped dramatically. This is likely due to 1) the AAB-AFP promoter had only half the transfection levels of CMV even in vitro in Hep3B cells and 2) the lower expression levels of p53-Bad* using the adenovirus. AAB-AFP may be more successful in expressing p53-Bad* in a different vector.

A main difference between our successful p53-Bad* in vitro and in vivo zebrafish experiments and the mixed results presented in this study is our use of the adenoviral vector as a delivery system. Though the expression levels of p53-Bad* in plasmid form were somewhat lower than that of WT p53^4^, the viral packaging may further exacerbate this issue, and we hypothesize that there is simply not enough p53-Bad* reaching the tumor to successfully induce cell death. The p53-Bad* construct is a toxic, fast acting construct that differs from WT p53 in terms of mechanism of action^4-5^. However, the construct still requires sufficient time for it to be expressed at an adequate level.

It is also likely that the starting size of the tumors had enough of an impact on the penetration of construct that repeating this study with a smaller overall and tighter limit on starting tumor size (e.g. 50-100 mm^3^), would reduce error from this source and demonstrate similar success between adenoviral p53-Bad* and WT p53. Increasing the frequency of dosage (to daily rather than 3x per week) and extending the study endpoint may also show more success for adenoviral p53-Bad*, as our in vitro data showed increased success for p53-Bad* compared with WT p53 as dosage increased and the timepoint was extended. Furthermore, previous data^15^ from our lab showed that pausing dosing of mitochondrially targeted p53 for more than one day allowed tumors to continue to grow. Thus, were this study to be repeated, the authors would suggest more frequent dosing of p53-Bad* for increased efficacy. This is not surprising given the short-acting nature of p53-Bad*. In zebrafish studies, p53-Bad* was not transiently expressed but integrated into the genome as a transgene^4^, circumventing transient transfection in that animal model.

In order to maximize success, it may be wise to further explore the reasons behind the significantly lower expression of p53-Bad* compared with WT p53 when delivered via adenovirus before initiating further animal studies. One possibility is that the mitochondrially-targeted p53-Bad* acts too quickly, reducing the amount of available cellular ATP enough to halt further transcription of p53-Bad* and allowing some of the cells to recover. Another is that, despite the CMV promoter being consistent across the constructs, the genes may be undergoing promoter activity buffering^16^, which has been shown to protect cells from misexpression of harmful genes. If the issues are specifically related to adenoviral delivery it may be best to package p53-Bad* in an alternative delivery vector such as a polymeric nanoparticle or AAV viral vector. Other options include increasing the number of copies of p53-Bad* in the current adenoviral vector, modifying the adenovirus to increase efficiency of delivery, or exploring CRISPR/Cas9 technologies for integration into the genome.

## Acknowledgements

We would like to gratefully acknowledge NIH Grant #R21CA241015-02 to Dr. Carol Lim.

Research reported in this publication utilized the Preclinical Research Shared Resource at Huntsman Cancer Institute at the University of Utah and was supported by the National Cancer Institute of the National Institutes of Health under Award Number P30CA042014. The content is solely the responsibility of the authors and does not necessarily represent the official views of the NIH.

## Supplemental Figures

**Figure S1:**
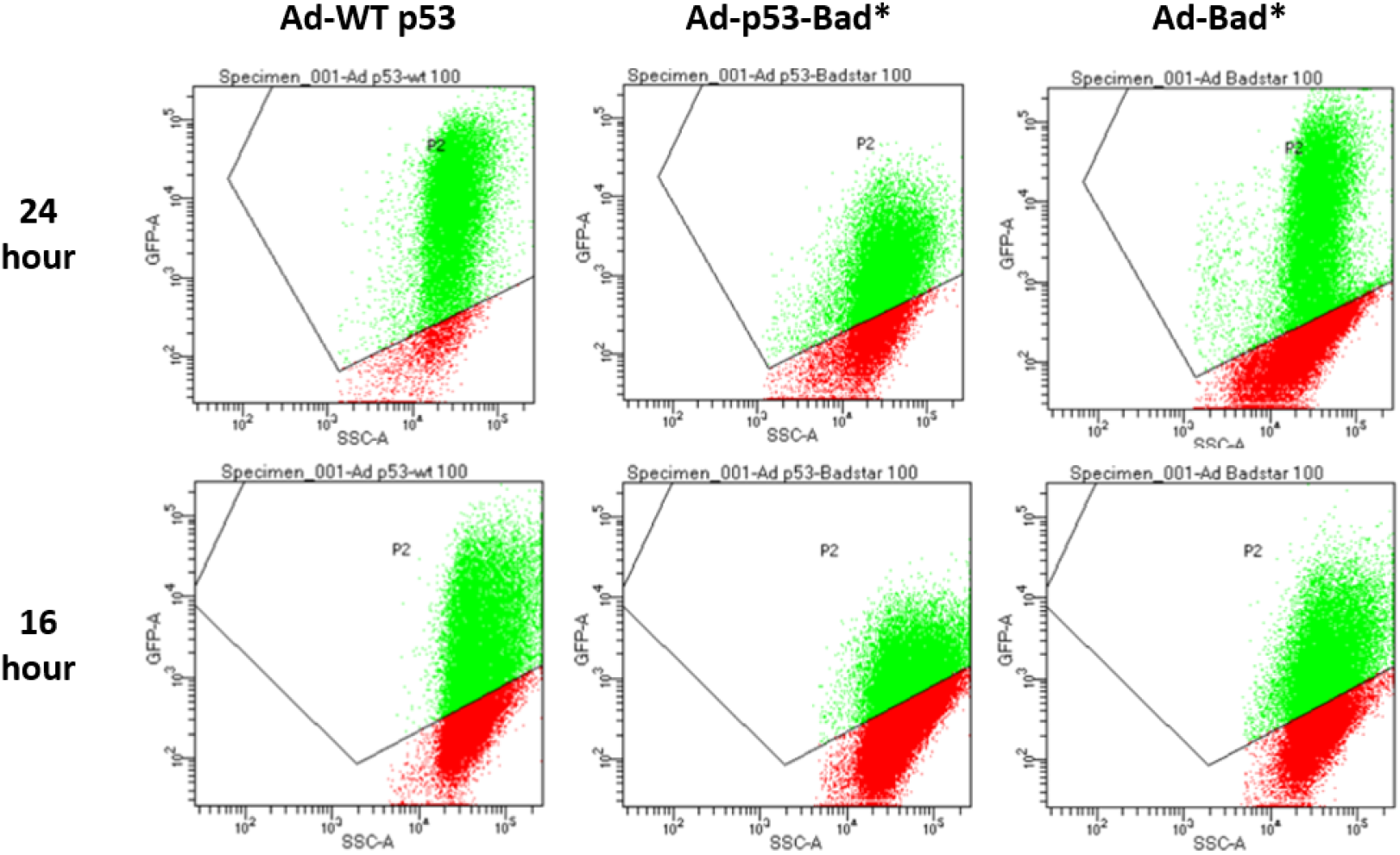
GFP expression levels. Adenoviral p53-Bad* treated cells showed distinctly lower expression compared with both adenoviral p53 and adenoviral Bad* at two different time points. Cells were gated for size (live cells only, no fragments) and GFP expression (indicated by P2 boundary). GFP-A (y axis) correlates to intensity of GFP-detection signal.

**Figure S2:**
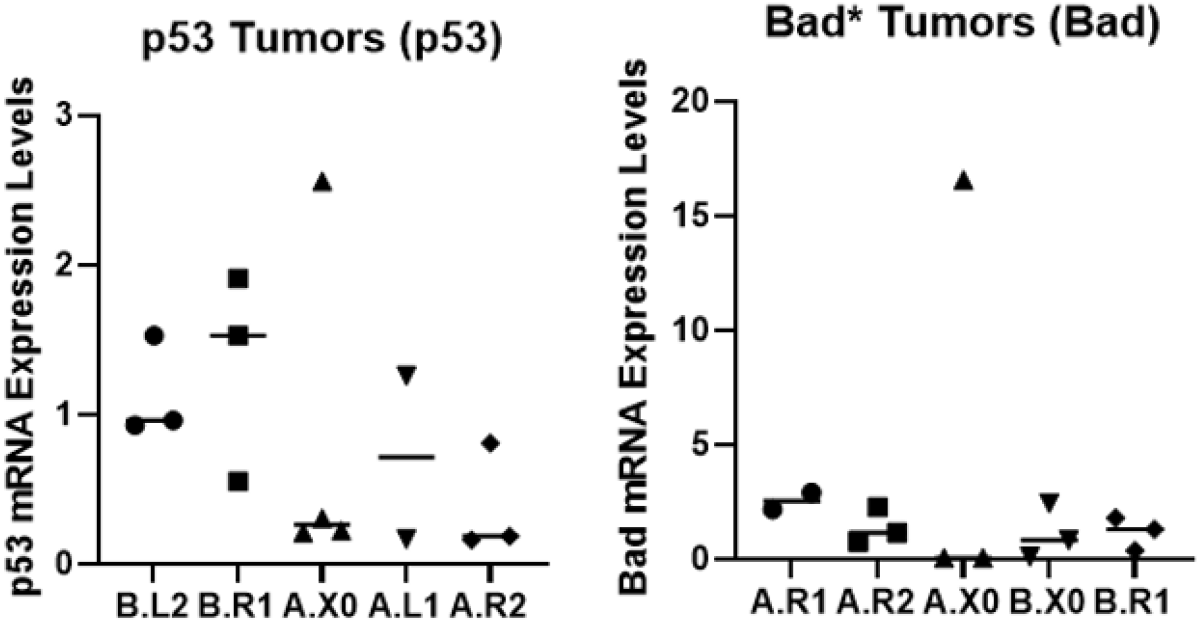
Data with outliers (before cleaning with Grubb’s test).

## Notes

### Competing Interest Statement

The authors have declared no competing interest.

